# The proteotoxicity of azetidine 2-carboxylic acid is associated with reactive oxygen species accumulation

**DOI:** 10.64898/2026.08.06.743286

**Authors:** K. M. Asha Alles, Devasantosh Mohanty, Varun Dwivedi, Ryo Yokoyama, Ron Mittler, Craig A. Schenck

**Author notes:** Corresponding author –.

## Abstract

Plants make diverse metabolites to outcompete neighboring organisms for space and resources. Some of these toxic metabolites broadly disrupt conserved molecular mechanisms, such as protein biosynthesis. Nonproteogenic amino acids (NPAAs) are a structurally diverse class of metabolites that interfere with protein biosynthesis. The proline (Pro) analog azetidine-2-carboxylic acid (Aze) inhibits plant growth through misincorporation during protein biosynthesis. However, it is unknown if a cascade of downstream stress responses is triggered following Aze misincorporation. Here, we investigate the morphological and stress responses in Arabidopsis grown on Aze. Investigation of root morphological responses show not only reduced root growth, but increased root branching following growth on Aze. Altered root morphology is coupled with a reduced gravitropic response. Aboveground organs were also affected by Aze, including reduced chlorophyll content, reduced photosynthetic efficiency, and increased anthocyanin content. We then tested whether Aze induces reactive oxygen species (ROS) accumulation using multiple approaches and observed both immediate and sustained accumulation of general ROS and H_2_O_2_ following treatment with Aze. When plants were grown on Aze supplemented with Pro, ROS levels were restored to normal levels, suggesting that reducing misincorporation events results in less downstream stress responses. In summary, we find that following Aze treatment a cascade of downstream stress responses is induced that exacerbates the effects of toxic NPAAs. This study sheds light on the mechanism of action of NPAAs and provides information on the downstream consequences of translational errors.

## Introduction

Plants make a vast repertoire of toxic metabolites and their mechanisms of action are as diverse as their chemical structures (Agrawal and Fishbein, 2006; Alseekh and Fernie, 2018; Schenck and Busta, 2021; Vendemiatti et al., 2024). Some are exquisitely specific for a single protein target in one organism, whereas others are broadly toxic by targeting conserved biological processes (Wink, 2015). Deploying a mixture of toxic compounds with varying mechanisms of action enables plants to outcompete neighboring organisms for space and resources (Agrawal and Fishbein, 2006; Endara et al., 2023). Broadly toxic metabolites that interfere with protein biosynthesis are good strategies to accomplish these goals.

One class of broadly toxic metabolites is nonproteogenic amino acids (NPAAs). NPAAs are structurally analogous to one of the 20 proteogenic amino acids, but are not typically incorporated during protein biosynthesis in the organisms that synthesize NPAAs (Jander et al., 2020). There are over 250 NPAAs reported across plants, however they appear to be enriched in some plant families like Fabaceae and Asparagaceae (Bell et al., 2008; Vranova et al., 2011; Gibson et al., 2025). Although structures and distribution of NPAAs are known in plants, their mechanisms of action and the cascade of stress responses that are triggered following NPAA treatment are less clear.

Azetidine 2-carboxylic acid (Aze) is a NPAA analog of proline (Pro). It is produced in restricted plant lineages including some legumes and monocots (Fowden, 1956; Sung and Fowden, 1969; Leete et al., 1974; Gibson et al., 2025). Aze inhibits the growth of a diverse range of plants as well as bacteria, insects, and human cell lines (Adeyeyé and Blum, 1989; Song et al., 2017; Biratsi et al., 2021; Thives Santos et al., 2024). Proteomics analysis following Arabidopsis growth on Aze found that Aze is misincorporated specifically at Pro positions at a global rate of around 6% (Thives Santos et al., 2024). However, misincorporation rates were variable for different proteins and positions within the protein. The unfolded protein response was enhanced following Aze treatment, suggesting that misincorporation results in misfolded proteins and translates into reduced growth (Thives Santos et al., 2024). However, the molecular responses following misincorporation events are not well characterized.

Here we use Aze as a tool to investigate the stress and morphological responses following NPAA treatment. We hypothesize that misincorporation triggers a cascade of stress responses that amplify the proteotoxic effects of Aze. To test this hypothesis, we grew Arabidopsis on Aze-containing media and characterized various stress responses. We find that root growth is reduced and branching increases following Aze treatment. Aze induces accumulation of stress metabolites including anthocyanins while reducing chlorophyll content and photosynthetic efficiency. Aze triggers rapid and sustained accumulation of ROS that can be recovered by addition of Pro, but not other antioxidants or osmoprotectants. Our data demonstrates that initial misincorporation events trigger a cascade of downstream stress responses that could potentially amplify the effects of Aze. Greater understanding of the molecular responses following amino acid misincorporation provides insight into the fidelity of protein translation and biological consequences when errors occur.

## Results

### Aze alters root phenotypes and gravitropic response

To define root responses to Aze, we grew Arabidopsis on various concentrations of Aze and monitored root growth by measuring the longest roots and all lateral roots. As Aze concentrations increased, longest root length was reduced (Fig. 1A), consistent with previous results (Lee et al., 2016; Thives Santos et al., 2024). At 10 µ M Aze root growth is inhibited by ∼80% and at higher concentrations, 100 µM, root length is almost completely inhibited (Fig. 1A). Consistently, the sum of the lengths of all roots was also decreased with increasing Aze concentrations, however the lateral root branching was increased (Fig. 1B &C). We analyzed the degree of root branching by calculating the ratio of the longest root over the total root length (Fig. 1C). If the length of the longest root and the total of all root length were the same, this indicates no root branching with a ratio of 1. As this ratio decreases it indicates more root branching. We found that at 10, 20, and 50 µM Aze the ratios were significantly reduced compared to growth on no Aze (Fig. 1C). However, at 100 µM Aze the ratios were not significantly different (Fig. 1C), likely due to the challenge of determining primary and lateral roots when growth is almost completely inhibited. These data show that Aze inhibits root growth and stimulates lateral root production.

**Fig. 1.**
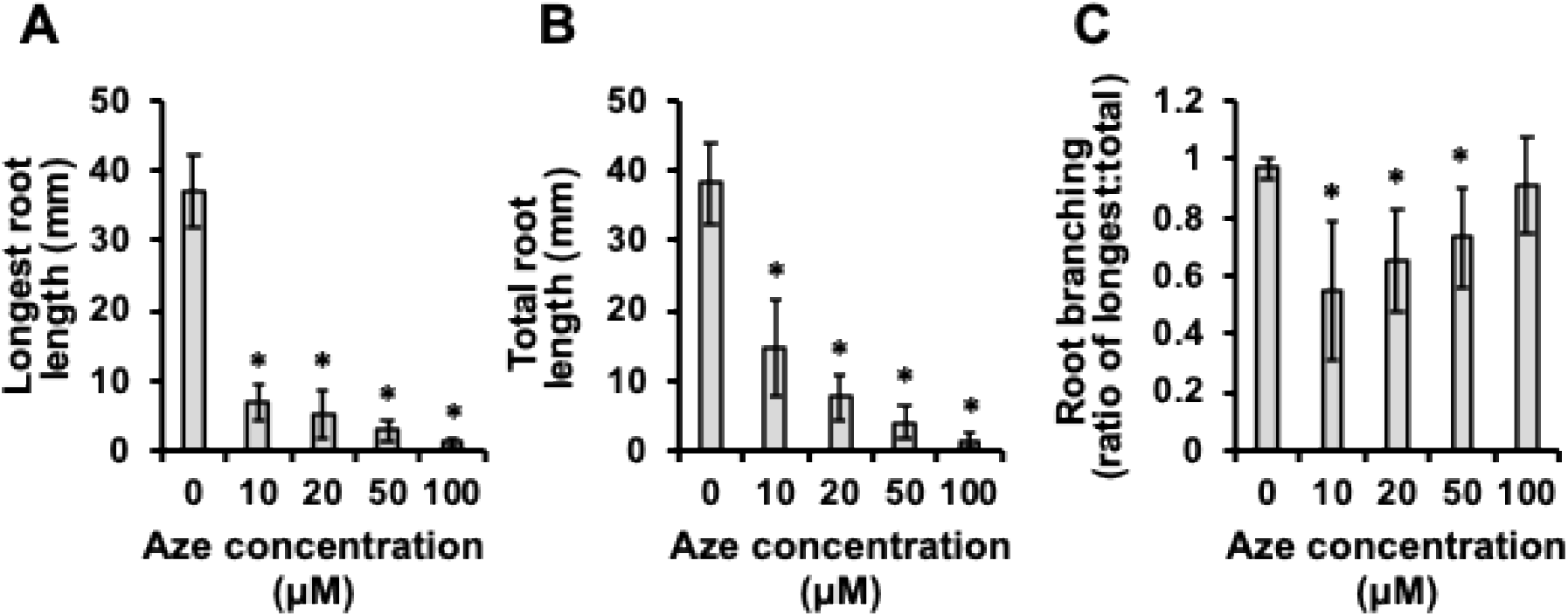
Aze inhibits Arabidopsis root growth and induces branching. Arabidopsis was grown on 0.5x MS media for 8 days supplemented with varying amounts of Aze. Root length was measured using ImageJ. **(A)** Length of the longest root, **(B) l**ength of all roots including the lateral roots, **(C)** ratio of the longest root to the total root length as a proxy for overall branching. A ratio of 1 indicates no branching, with lower values indicating higher root branching. Bars indicate mean ± SEM of ≥ 10 biological replicates. Stars indicate significant differences from 0 μM Aze control in a one-way T-test of P < 0.05. Abbreviation: Aze, Azetidine 2-carboxylic acid.

Given the enhanced lateral root formation, we hypothesized that Aze treatment might alter auxin concentrations in the roots. To test this hypothesis, we measured the gravitropic response of roots grown on Aze. Since high Aze concentrations severely reduced root length, we grew plants on 5 µM Aze so that roots were long enough to monitor a gravitropic response, however still effected by Aze. Arabidopsis seedlings were grown for 8 days on 0 or 5 µM Aze then rotated 90 degrees, and the angle of curvature was measured after 72 hours (Supplementary Fig. 1A). A 90 degree angle represents a full gravitropic response, whereas values greater than 90 represent a reduced gravitropic response. Arabidopsis grown on 0 µM Aze almost reached a 90 degree angle following reorientation, indicating a full gravitropic response, however the response of plants grown on 5 µM was significantly reduced, only reaching an angle of 120 degrees on average (Supplementary Fig. 1B). We attempted to test if higher Aze concentrations had a more severe impact on gravitropic response, however at higher concentrations, root growth was too severely reduced that reorientation angles were unable to be determined accurately.

### Aze induces above ground stress responses

Although Aze has dramatic effects on root growth phenotypes, we wanted to test the effects of Aze on above ground phenotypes. We grew plants on Aze and measured chlorophyll content, photosynthetic efficiency, and anthocyanin content. Total chlorophyll content was measured from seedlings following growth on 20 and 100 µM Aze for 8 days (Fig. 2A). Total chlorophyll content was reduced at 100 µM Aze, however 20 µM did not reduce total chlorophyll content compared to seedlings grown on 0 µM Aze (Fig. 2A). Individual chlorophyll a and b showed a similar trend to total chlorophyll, reduced at 100 µM, however not significantly different at 20 µM (Supplementary Fig. 2A). Given the reduction in chlorophyll content, we hypothesized that photosynthetic capacity would be reduced during Aze treatment. To test this, we measured maximum quantum yield of Photosystem II (Fv/Fm) from Arabidopsis grown on 0 and 100 µM Aze (Murchie and Lawson, 2013). Following growth on Aze, we found that Fv/Fm was reduced in old leaves. However newly emerged leaves from plants grown on Aze were not significantly different from new leaves on control plates (Fig. 2B and Supplementary Fig. 2B). Next, we measured anthocyanin content from leaves, another key indicator of stress response (Kovinich et al., 2015). Accumulation of purple compounds on the undersides of leaves following treatment with high Aze concentrations was observed (Supplementary Fig. 2C), thus we hypothesized that this was due to anthocyanin accumulation. Anthocyanins were measured spectrophotometrically following growth for 8 days on 20 and 100 µM Aze. We found that anthocyanin content was significantly increased following Aze treatment at both 20 µM and 100 µM (Fig. 2C). These data show that Aze affects both above and below ground growth and stress responses.

**Fig. 2.**
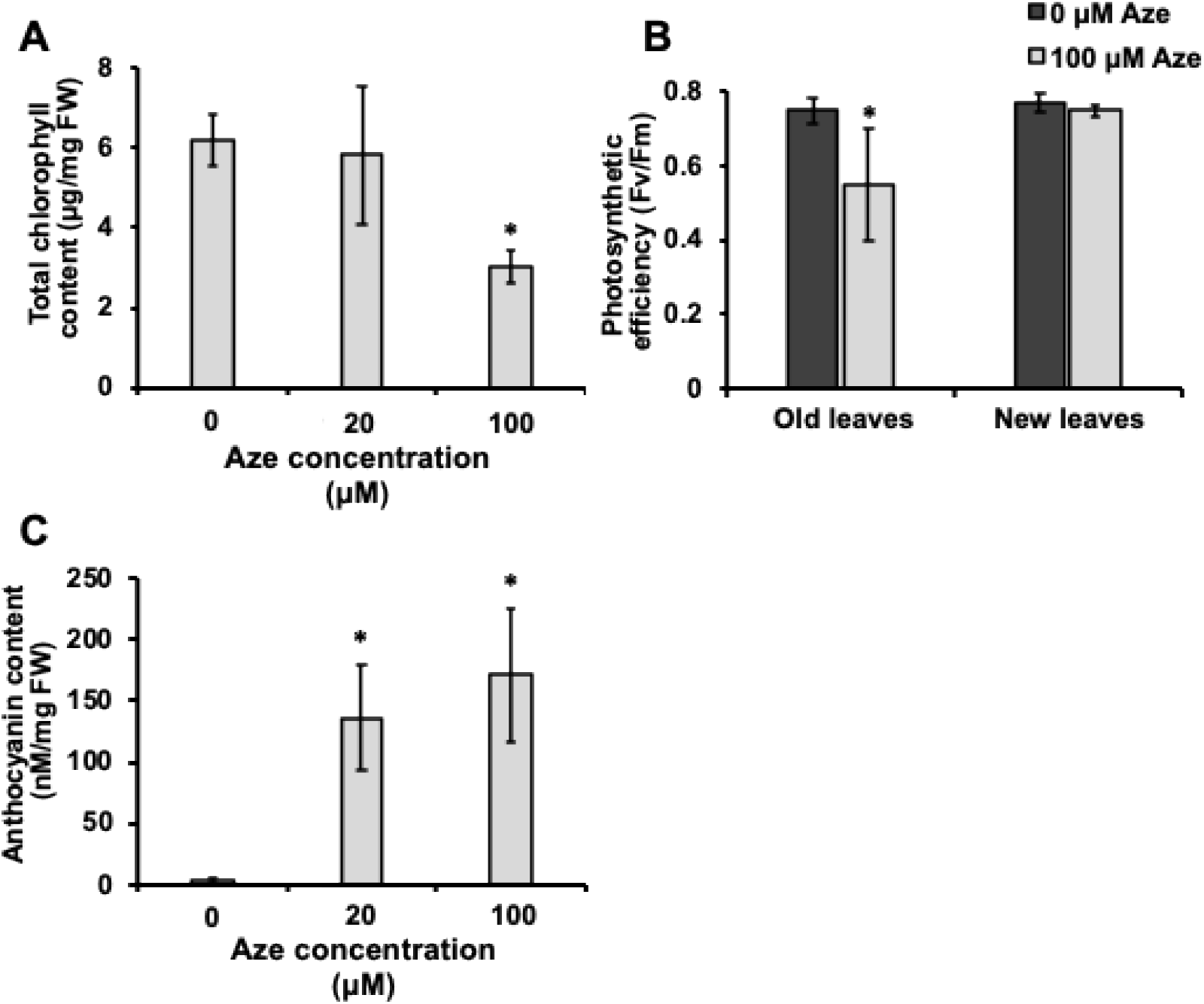
Aze triggers above ground stress phenotypes. Arabidopsis was grown on 0.5x MS media for 8 days supplemented with varying amounts of Aze. Above ground stress phenotypes were measured: **(A)** chlorophyll content, **(B)** photosynthetic efficiency, and **(C)** anthocyanin content. Total chlorophyll content is shown in **(A)** and was consistent with differences observed in chlorophyll a and b (Supplementary Fig. 2A). **(B)** The Fv/Fm levels in old and young leaves of Arabidopsis treated with 0 or 100 µM Aze. A representative image of Fv/Fm can be found in Supplementary Fig. 2. **(C)** Anthocyanin content. Bars indicate mean ± SEM of ≥ 10 biological replicates. Stars indicate significant differences from 0 μM Aze control in a one-way T-test of P < 0.05. Abbreviations: Aze: Azetidine 2-carboxylic acid, FW: Fresh weight, Fv/Fm: maximum quantum efficiency of photosystem II.

### Aze induces reactive oxygen species accumulation

To test if reactive oxygen species (ROS) accumulate following Aze treatment, we first grew plants for 8 days on increasing Aze concentrations and measured H_2_O_2_ levels using the Amplex-Red (10-acetyl-3,7-dihydroxyphenoxazine) method from whole seedlings (Fig. 3A). H_2_O_2_ levels were significantly increased at 50 and 100 µM Aze concentrations, however lower Aze concentrations were not significantly different from control (Fig. 3A). We next wanted to measure more general ROS accumulation in plant tissues. To test this, we grew plants on 10 and 20 µM Aze for 8 days, and measured ROS using a ROS reactive fluorescent dye, dichloroflourescein (H_2_DCFDA), that measures intracellular ROS accumulation (Fichman et al., 2019; Fichman et al., 2022; Mohanty et al., 2026a). At higher concentrations of Aze (50 and 100 µM) plant growth was too severely reduced that it was challenging to accurately determine ROS accumulation from the images. However, at 10 µM Aze, ROS accumulation was significantly increased compared with controls (Fig. 3B), similar to increased H_2_O_2_ levels a high Aze concentrations (Fig. 3A). Although we observe increase in ROS accumulation following 8 days of growth on Aze, we wanted to test if ROS accumulate rapidly following Aze treatment. To test this, we grew plants on control media for 6 days, then fumigated plants with either a buffer control or 100 µM Aze, both containing H_2_DCFDA, and measured ROS accumulation after 30 minutes (Fig. 3C). Here, given the long roots we can observe organ-specific accumulation of ROS (Fig. 3C). We observe a significant increase in intracellular ROS accumulation following Aze that appears to be specific to the roots (Fig. 3C). Quantification of DCF intensity shows a significantly increased rapid accumulation of general ROS following a rapid 30 minute Aze treatment (Fig. 3D). These data suggest that both rapid and sustained ROS accumulation are another consequence of Aze treatment.

**Fig. 3.**
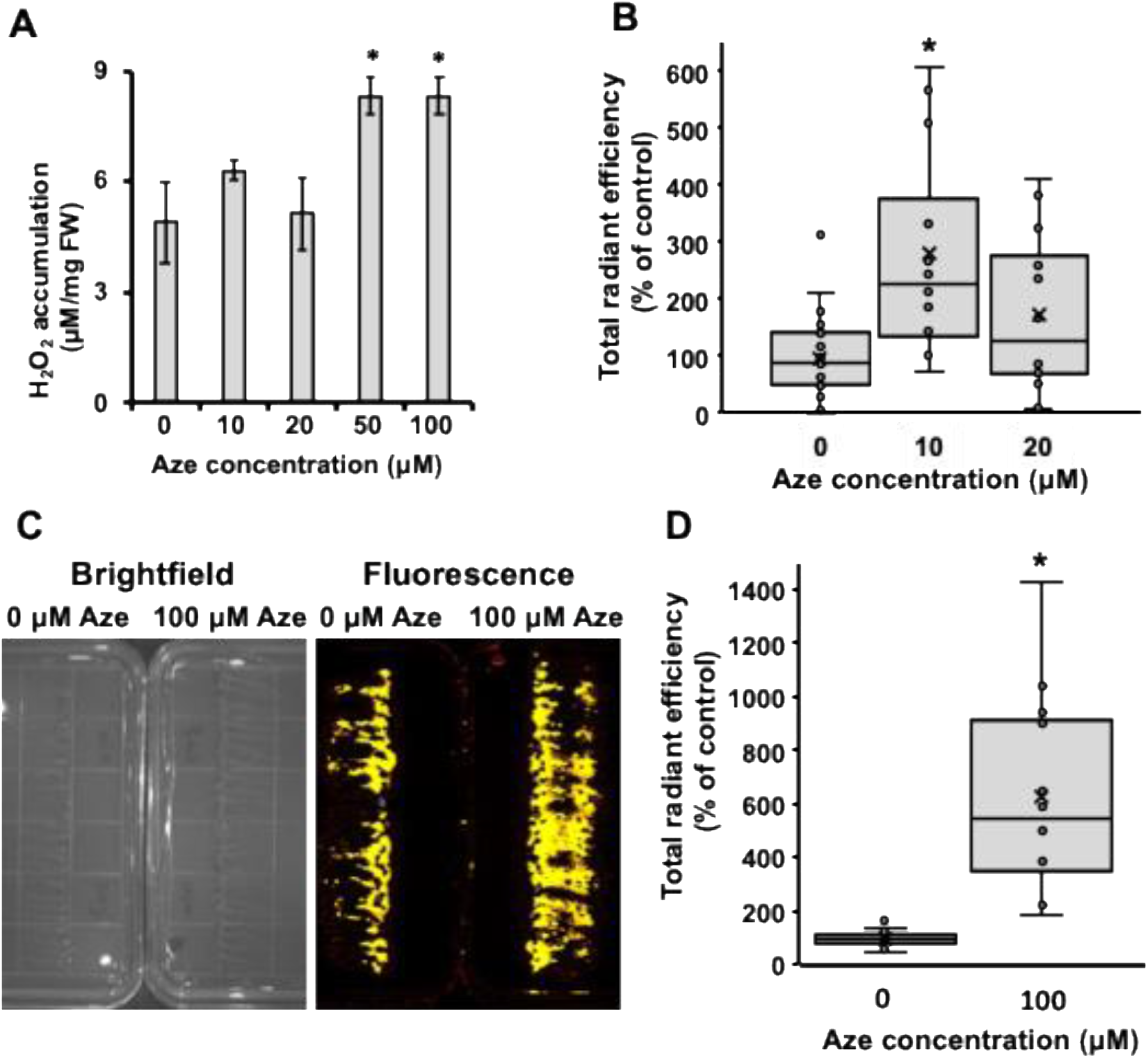
Aze triggers rapid and sustained reactive oxygen species (ROS) accumulation. Arabidopsis seedlings were grown on 0.5x MS media for 8 days supplemented with varying amounts of Aze (**A**,**B**). **(A)** H_2_O_2_ levels were measured from total seedlings grown on different concentrations of Aze using an Amplex Red assay. Significantly increased H_2_O_2_ levels were observed at high Aze concentrations. Bars indicate mean ± SEM of ≥ 10 biological replicates. Stars indicate significant differences from 0 μM Aze control in a one-way T-test of P < 0.05. **(B)** General intracellular ROS were measured from total seedlings grown on different concentrations of Aze using a fluorescent ROS reactive stain (H_2_DCFDA). Intensity of signal was measured using an IVIS imager and normalized to 0 μM controls. Plots show mean and all data points collected. Stars indicate significant differences from 0 μM Aze control in a one-way T-test of P < 0.05. **(C**,**D)** Arabidopsis seedlings were grown for 8 days on control media, then fumigated with a buffer control or 100 μM Aze (both containing H_2_DCFDA) and intracellular ROS was measured using an IVIS imager after 30 minutes. **(C)** Representative image showing root-specific accumulation of ROS following Aze treatment. **(D)** Quantification of ROS intensity using ImageJ. Plots show mean and all data points collected. Stars indicate significant differences from 0 μM Aze control in a one-way T-test of P < 0.05. Abbreviations: Aze: Azetidine 2-carboxylic acid, FW: Fresh weight.

### Antioxidants do not restore Aze-induced root growth reduction

Given the accumulation of ROS following Aze treatment, we hypothesized that root growth could be restored by addition of antioxidants similar to the restoration of root growth by adding Pro (Thives Santos et al., 2024). To test this hypothesis, we grew plants on Aze plus various antioxidants, including mannitol, sorbitol, glutathione, ascorbic acid, N-acetyl-L-cysteine, and L-cysteine, and monitored root growth. Previously, we found that between a 5-10-fold excess of Pro was sufficient to fully restore root growth of Arabidopsis on Aze (Thives Santos et al., 2024). Thus, we grew Arabidopsis on 20 µM Aze with 200 µM of various antioxidant compounds. As a control for restored growth, we also included a 10-fold excess of L-Pro (Supplementary Fig. 3). In this experiment, Aze reduced root growth by ∼87% and addition of L-Pro was sufficient to fully restore growth to control levels, indicating that L-Pro could mitigate the misincorporation effects of Aze on root growth (Supplementary Fig. 3). However, none of the antioxidants tested had any significant impacts on restoring root growth (Supplementary Fig. 3). These data could suggest that accumulation of ROS is a downstream consequence of Aze treatment, and application of compounds that could act to scavenge ROS is not sufficient to restore the initial events of Aze misincorporation into proteins.

### Pro restoration of ROS accumulation

Although Pro restores growth following Aze treatment, we wanted to test if Pro also restored ROS accumulation to normal levels. To test this hypothesis, we performed two experiments to test the rapid and sustained ROS accumulation following Aze treatment. First, we grew plants on control media, media with 100 µM Aze or media with 100 µM Aze plus 1000 µM Pro for 8 days. Seedlings were harvested and used for total H_2_O_2_ levels. Similar to our previous results of plants grown on 100 µM Aze containing media, H_2_O_2_ accumulation was observed following long-term Aze exposure compared with the control (Fig. 4A). However, when plants were grown on media containing Aze and Pro, H_2_O_2_ levels were significantly different from Aze alone and similar to the controls (Fig. 4A), suggesting that Pro restored normal ROS levels. Next, to test the rapid response, we grew plants for 7 days on 0.5x MS control plates. Then, we transferred the seedlings to well plates with water. After 1 day of recovery, water was replaced with either fresh water, 100 µM Aze or 100 µM Aze plus 1000 µM Pro. Total H_2_O_2_ levels were measured 60 minutes following replacement. Growth in well plates enabled us to rapidly change treatments while minimizing physical stress-induced ROS accumulation by moving plants from plate to plate. Similar to our previous results of plants exposed to 100 µM Aze for 30 minutes, H_2_O_2_ accumulation was observed following short-term Aze exposure compared with the control (Fig. 4B). However, when plants were exposed to Aze and Pro, H_2_O_2_ levels were not significantly different from the control and significantly reduced compared to Aze alone (Fig. 4B), suggesting that Pro restored normal ROS levels.

**Fig. 4.**
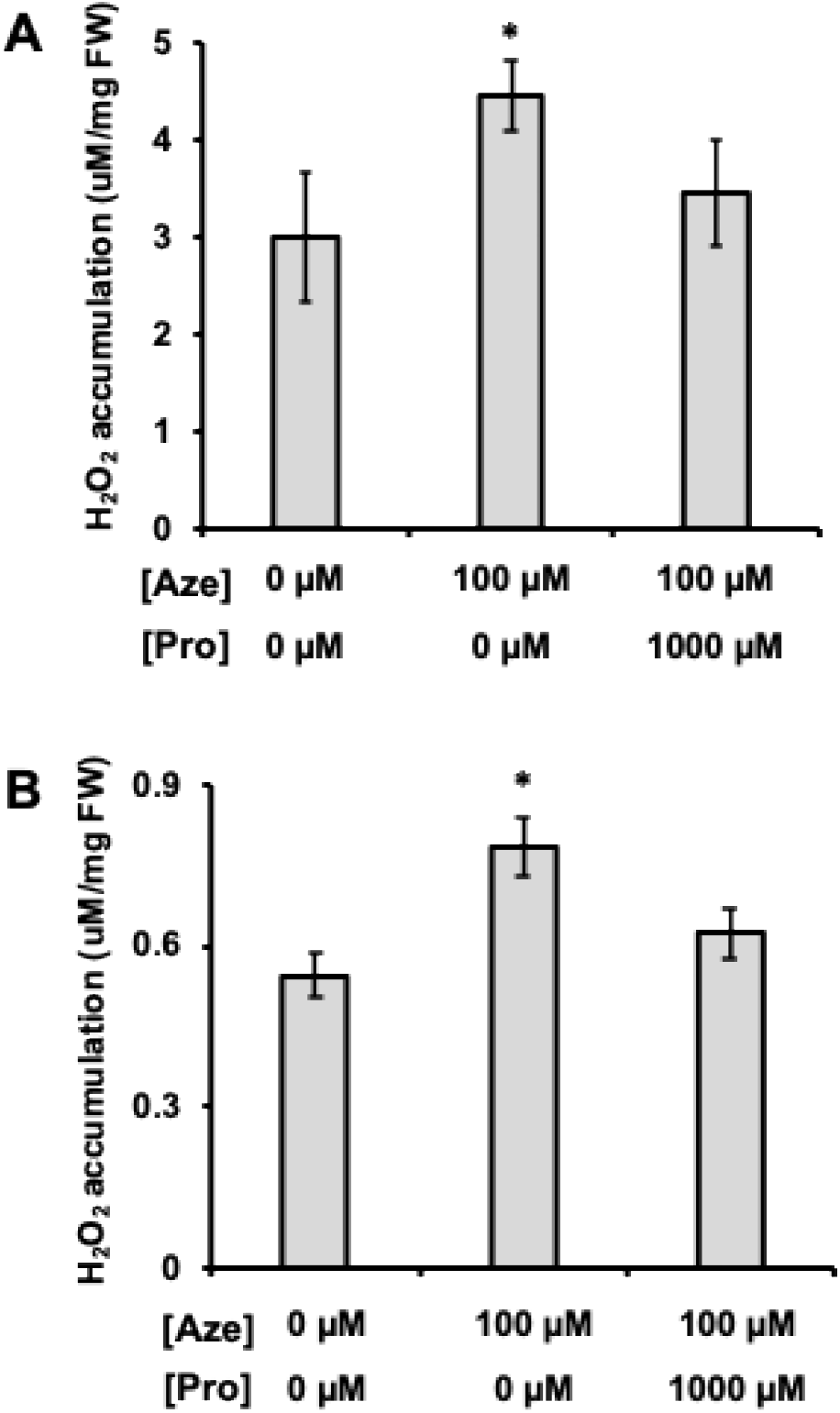
Proline restores H_2_O_2_ accumulation. Arabidopsis was grown in the presence of Aze alone or Aze plus proline (Pro). H_2_O_2_ levels were measured using an Amplex red kit after 8 days of treatment **(A)** and after 60 minutes of treatment (**B**). **(A)** Arabidopsis was grown for 8 days and then total seedlings were used for H_2_O_2_ measurements. H_2_O_2_ levels were restored to control levels when the Aze media was supplemented with a 10-fold excess of proline. (**B**) Arabidopsis was grown for 6 days in control liquid media in well plates. Then, the media was removed and replaced with control media, media containing 100 μM Aze alone, or media containing 100 μM Aze plus 1000 μM proline. H_2_O_2_ levels were measured whole seedlings after 60 minutes of treatment. Bars indicate mean ± SEM of ≥ 10 biological replicates. Stars indicate significant differences from 0 μM Aze control in a one-way T-test of P < 0.05. Abbreviations: Aze: Azetidine 2-carboxylic acid, FW: Fresh weight.

### Aze induces ROS responsive genes

ROS accumulation is induced following biotic and abiotic stress responses as well as Aze (Mittler et al., 2022; Peláez-Vico et al., 2024). Various transcription factors and ROS related genes have been shown to increase in their expression following excess-light stress treatments that induce ROS (Fichman et al., 2022; Fichman et al., 2024). Thus, we tested whether previously implicated transcripts involved in ROS accumulation were induced following Aze treatment. Plants were grown for 5 days on 0.5x MS control solid media plates and moved to either control media or media containing 100 µM Aze. RNA was extracted from whole seedlings 1, 3 and 6 hours after moving onto new media. In general, the expression of ROS inducible transcripts was increased following Aze treatment (Fig. 5), consistent with previous studies (Fichman et al., 2022; Fichman et al., 2024). Of the five transcripts monitored, four showed significantly increased expression in at least one time point during Aze treatment (Fig. 5). *AtAPX2* showed the highest induction of about 200-fold after 1 hour, subsiding to 50-fold after 6 hours on Aze, compared to the control (Fig. 5). *AtZAT10* followed a similar expression pattern as *AtAPX2* but was elevated to a lesser extent, around 4-fold at each time point (Fig. 5). *AtZAT12* was significantly increased following Aze treatment at 1 and 6 hours, but reduced at 3 hours, while *AtRBOHD* was significantly enhanced only at 3 and 6 hours (Fig. 5). *AtMYB30* was the only transcript that showed no change in expression following Aze treatment compared with controls (Fig. 5). These data indicate that in general ROS related transcripts are induced, however Aze might stimulate a specific ROS response rather than generally turning on all ROS related genes.

**Fig. 5.**
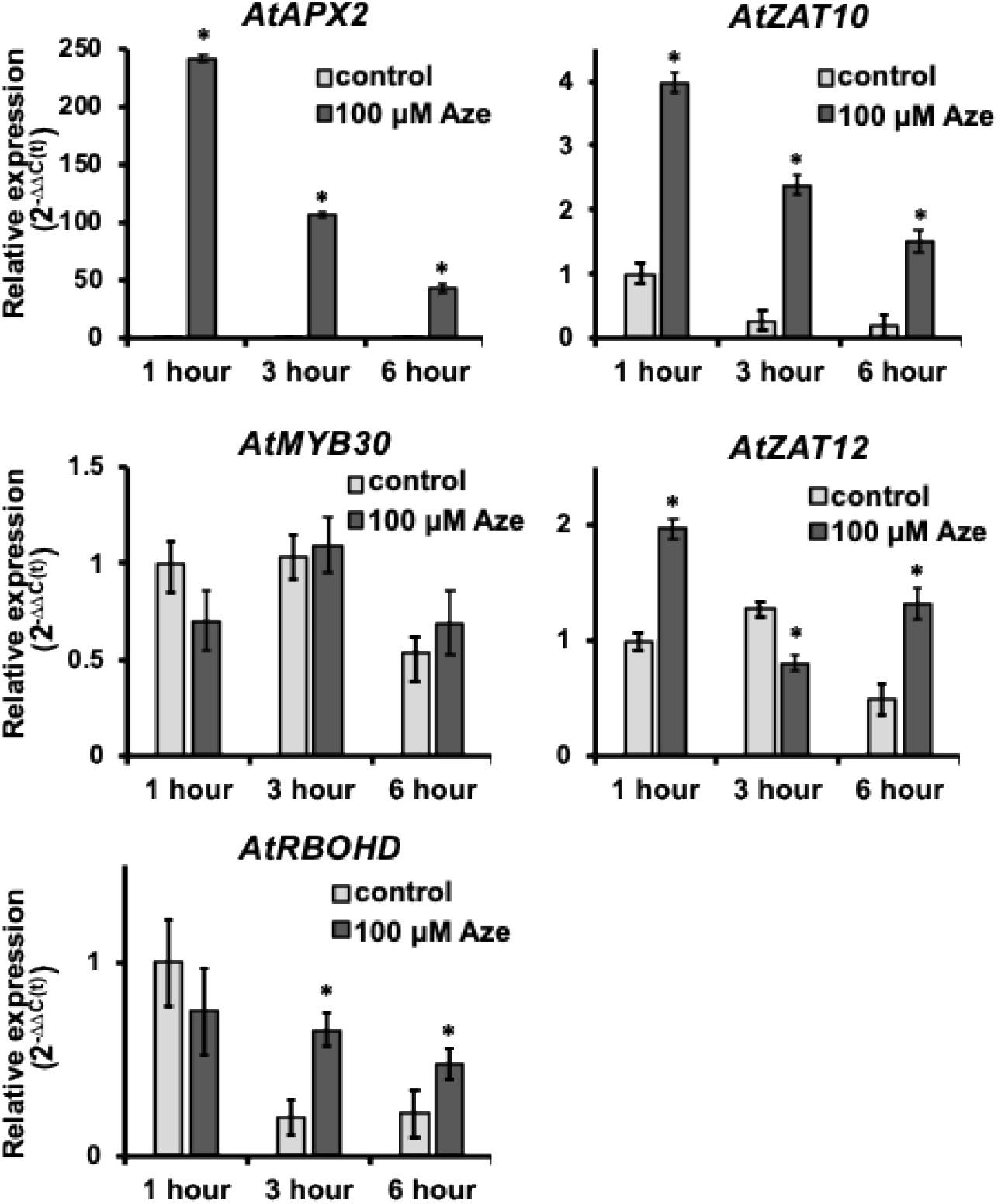
Aze induces expression of ROS related genes. Expression of ROS related genes was measured following 1-6 hours of Aze treatment. Arabidopsis seedlings were grown for 5 days on control solid media, then moved to control media or media containing 100 μM Aze. RNA was extracted from seedlings after 1, 3, and 6 hours of treatment. ROS related genes were measured using qRT-PCR, and relate expression was quantified using 2^-ΔΔC(t)^ in comparison to a housekeeping gene. Bars indicate mean ± SEM of ≥ 3 biological replicates. Stars indicate significant differences from 0 μM Aze control at each time point in a one-way T-test of P < 0.05. Abbreviations: APX2: Ascorbate peroxidase 2, AZE: Azetidine 2-carboxylic acid, MYB30: Myeloblastosis domain protein 30, RBOHD: respiratory burst oxidase homolog D, ZAT10: Zinc finger of *Arabidopsis thaliana* 10, ZAT12: Zinc finger of *Arabidopsis thaliana* 12.

## Discussion

Non-proteogenic amino acids are prevalent across plants and some are used as a generalist defense strategy to outcompete neighboring organisms (Bell, 2003; Bertin et al., 2007; Huang et al., 2011; Jander et al., 2020). A recent study found that Aze is misincorporated in place of Pro across the whole proteome in Arabidopsis resulting in reduced root growth demonstrating that Aze misincorporation is the major mechanism of action (Thives Santos et al., 2024). We hypothesized that Aze misincorporation leads to protein misfolding and aggregation that triggers a more global stress response. Here, we directly tested the hypothesis that Aze misincorporation triggers a cascade of downstream stress responses.

Aze has similar stress induced responses as other NPAAs. For example, meta-tyrosine, canavanine, and theanine all inhibit root growth and alter lateral root branching (Krasuska et al., 2016; Zer et al., 2020; Chen et al., 2023). Growth of Arabidopsis on Aze-containing media alters both above and below-ground phenotypes. Below ground, root growth is inhibited, but branching is induced by Aze (Fig. 1). Additionally, the gravitropic response is attenuated by growth on Aze (Supplementary Fig. 1). These root phenotypes are likely associated with altered auxin maxima and minima across the roots, however it is not clear how Aze alters auxin concentrations in the roots (Roychoudhry and Kepinski, 2022). Aboveground, various stress responses were observed, including loss of chlorophyll content and reduced photosynthetic efficiency as well as accumulation of anthocyanin stress pigments (Fig. 2). Aze likely does not specifically inhibit chlorophyll biosynthesis, but reduced chlorophyll is a side-effect of misincorporation and protein misfolding and turnover (Li et al., 2025). This signals a stress response and thus why anthocyanins are also accumulated during Aze treatment. Alternatively, Aze may block the production and therefore the assembly of light-harvesting chlorophyll binding complexes (LHC) and thereby reduce chlorophyll levels (Levin and Schuster, 2023).

Another common response to stress is the accumulation of ROS (Mittler et al., 2022). We found that Aze induces both rapid and sustained ROS accumulation using two different approaches (Fig. 3). Interestingly, Aze-induced root growth could not be restored by supplementing antioxidants that might scavenge ROS but is restored only by Pro (Supplementary Fig. 3). Furthermore, growth on Aze and Pro alleviated the ROS accumulation observed when Arabidopsis was grown on Aze alone (Fig. 4). These data suggest that ROS accumulation is downstream of Aze misincorporation during protein biosynthesis and is another stress side-effect or stress-related signal. ROS imaging using H_2_DCFDA enabled the tissue-specific intracellular investigation of ROS accumulation. It appears that Aze-derived ROS are specifically accumulated in the roots (Fig. 3C). A previous study found that the unfolded protein response was more enriched in the roots compared with the leaves following Aze treatment, suggesting that the roots might experience greater stress in the presence of Aze resulting in more ROS accumulation in the roots (Thives Santos et al., 2024). ROS accumulation is also observed in response to treatment of other NPAAs (Konovalova et al., 2015; Krasuska et al., 2016; Chen et al., 2023), thus this is likely a conserved stress response following NPAA misincorporation.

This study provides greater insight into the mechanism of action of the NPAA Aze. Our findings suggest that the main cause of all downstream stresses is the initial misincorporation events of Aze that likely result in misfolding and nonfunctional proteins. Thus, general stress responses are turned on, such as ROS and anthocyanin accumulation, as well as reduced photosynthesis. Although NPAAs are structurally diverse, many appear to share a similar mechanism of action through misincorporation, followed by downstream stress responses such as accumulation of ROS and reduced photosynthetic capacity (Bertin et al., 2007; Konovalova et al., 2015; Krasuska et al., 2016; Zer et al., 2020; Chen et al., 2023). As a general defense strategy misincorporation during protein biosynthesis is a robust mechanism and evolving a tolerance mechanism is likely challenging. Although there are a few examples of bacteria growing in close association with the roots of Aze-producers that have evolved Aze detoxification enzymes (Nomura et al., 2003; Gross et al., 2008). This study provides molecular insight into the generalist proteotoxic stress responses of Aze and can be applied more broadly to the large class of plant-derived NPAAs.

## Materials and methods

### Plant materials, growth conditions, and Aze treatments

*Arabidopsis thaliana* ecotype Col-0 was used as plant material. Plants were grown on 0.5x MS (Murashige and Skoog) basal medium agar plates (0.6%) with Vitamins (PhytoTech Labs, Lenexa, KS, USA) and 1% sucrose. The media was supplemented with L-azetidine-2-carboxylic acid (Aze, Sigma) of varying concentrations. Seeds were surface sterilized with chlorine gas (80 mL bleach + 20 mL 12 M hydrochloric acid) for 1 hour. Sterilized seeds were placed on the media in square petri dishes (100 x 100 mm). Before positioning plates under the lights, they were placed at 4°C for 24 h. Plants were grown under controlled growth conditions: 23°C, 16-h daylight and 8-h dark photoperiod, and light intensity was ∼6800 lux. Plants were grown vertically for all the analyses unless stated otherwise in the figure legend.

### Root phenotype analysis

Plants were grown vertically for 8 days with different concentrations of Aze (0 µM, 10 µM, 20 µM, 50 µM, 100 µM). Plates were photographed, and root lengths were measured using ImageJ software. Both the longest root length and the total root length were measured, and the ratio of the longest root length to total root length was calculated.

### Gravitropic response analysis

Plants were grown vertically for 8 days on 0.5x MS medium containing either 0 or 5 µM Aze. Plates were subsequently rotated 90°, and root growth was permitted to proceed for an additional 3 days. The angle formed between the newly elongated root and the pre-existing root axis was then measured using ImageJ software.

### Chlorophyll measurement

Arabidopsis seedlings were grown for 8 days on 0.5x MS medium supplemented with 0, 20, or 100 µM Aze. For chlorophyll extraction, three seedlings were transferred to a 1.5 mL microcentrifuge tube containing 1 mL of 80% acetone and gently rocked at room temperature for 24 hours in the dark. Samples were then centrifuged at 15,000 x g for 7 minutes, and the resulting supernatant was used for spectrophotometric analysis. Absorbance was measured at 646 nm and 663 nm and chlorophyll content was determined from absorbance values as described in Lichtenthaler, 1987.

### Anthocyanin measurement

Arabidopsis seedlings were grown for 8 days on 0.5x MS medium supplemented with 0, 20, or 100 µM Aze. For anthocyanin extraction, three seedlings per sample were transferred to a microcentrifuge tube and homogenized thoroughly. The homogenate was resuspended in 400 µL of MeOH/CHCl_3_ (2:1 v/v), followed by the addition of 300 µL of H_2_O and 125 µL of CHCl_3_.

Samples were vortexed thoroughly and centrifuged at 15,000 × g for 10 minutes at room temperature to facilitate phase separation. The upper aqueous/methanolic phase, containing the anthocyanins, was carefully collected. To acidify and stabilize the anthocyanin-containing fraction, an equal volume of 0.1M HCl (300 µL) was added to 300 µL of the collected phase. Absorbance of the resulting solution was measured spectrophotometrically at 520 nm similar to Schenck et al., 2020.

### Photosynthesis measurement

Arabidopsis seedlings were grown on 0.5x MS medium for 8 days. Then they were transferred to new 0.5x MS media supplemented with 0 and 100 µM Aze and grown for 3 days. After at least 20 min dark adaptation, maximum quantum efficiency of photosystem II (Fv/Fm) was quantified using HEXAGON-IMAGING-PAM (Walz).

### H_2_O_2_ measurement

Arabidopsis seedlings were grown for 8 days on 0.5x MS medium supplemented with 0, 10, 20, 50, and 100 µM Aze. Hydrogen peroxide (H_2_O_2_) content was determined using the Amplex Red Hydrogen Peroxide Assay Kit (Invitrogen) following the protocol described by Brumbarova et al., 2016. Briefly, 100 mg of whole seedling tissue per treatment was harvested, immediately frozen in liquid nitrogen, and ground to a fine powder. Tissue extracts were prepared by resuspending the powder in 500 µL of 0.25 M sodium phosphate buffer (pH 7.4). H_2_O_2_ concentration was subsequently determined from the extract corresponding to 1 mg of fresh plant tissue per reaction, as outlined in the manufacturer’s protocol.

### ROS assay – using H_2_DCFDA

Arabidopsis seedlings grown on 0.5x MS media with different concentrations of Aze. The seedlings were grown for 8 days on 0.5x MS medium supplemented with Aze at concentrations of 0, 10, 20, 50, and 100 µM. Following the growth period, ROS were visualized *in planta* by fumigating the plates with 2′,7′-dichlorofluorescein diacetate (H_2_DCFDA) for 30 minutes. Post fumigation, images were captured using the IVIS Lumina S5 platform using Living Image 4.7.3 software in acquisition mode (PerkinElmer, Waltham, MA, USA) for 30 minutes. Images were processed using the Living Image 4.7.3 software (PerkinElmer) as described by Fichman et al., 2019, 2022 and the raw integrated density of regions of interests (ROIs) were calculated using Image J (Mohanty et al., 2026a; Mohanty et al., 2026b). In a complementary approach, Arabidopsis seedlings grown for 6-7 days on 0.5x MS medium were co-fumigated with 100 µM Aze and DCF-DA or DCF-DA buffer control and images were captured and analyzed after 30 minutes as described above.

### Antioxidant test

Arabidopsis seedlings were grown on 0.5x MS medium supplemented with Aze supplemented with different antioxidants. Here, we used 20 uM Aze + 200 uM of each antioxidant. We used L-Pro, mannitol, sorbitol, glutathione, ascorbic acid, N-acetyl-L-cysteine, and L-cysteine (Sigma). Unsupplemented 0.5x MS medium served as the negative control. Root length was measured using imageJ after 8 days of growth.

### H_2_O_2_ measurement in plants grown with Aze + Pro

To assess H_2_O_2_ accumulation under Aze with Pro exposure, Arabidopsis seedlings were grown for 8 days on 0.5x MS medium supplemented with 0 µM Aze, 100 µM Aze, and 100 µM Aze in combination with 1000 µM Pro. Following the growth period, H_2_O_2_ concentrations were quantified using the Amplex Red Assay Kit as described above.

To assess the rapid H_2_O_2_ response, 7-day-old seedlings grown on 0.5x MS medium were transferred to a multi-well plate containing water and placed on an orbital shaker for 24 hours to allow equilibration. Following equilibration, the water was removed by pipetting and replaced with either water, 100 µM Aze, or 100 µM Aze supplemented with 1000 µM Pro. H_2_O_2_ concentrations were measured after 1 hour of treatment using the Amplex Red assay.

### Gene expression analysis of ROS related genes

Quantitative RT-PCR (qRT-PCR) was used for gene expression analysis for ROS related genes as performed previously (Fichman et al., 2024; Peláez-Vico et al., 2024). Plants were grown on control media for 5 days and then moved to plates containing 0 or 100 µM Aze and total RNA was extracted from seedlings at 1, 3, and 6 h after transfer to the new medium using the RNeasy Plant Mini Kit (QIAGEN). First-strand cDNA was synthesized using All-In-One 5× RT MasterMix (Applied Biological Materials) and diluted 5-fold to 10-fold before analysis. Amplification efficiency was determined for each primer pair. Each 10-µL RT–qPCR reaction contained 5 µL of Fast SYBR Green Master Mix (Applied Biosystems), 200 nM of each gene-specific primer, and diluted cDNA. *AtEF-1α* was used as the reference gene for normalization. RT–qPCR was performed as described previously (Thives Santos et al., 2024). The primer sequences used in this study are listed in Supplemental Table 1.

## Acknowledgments

We thank the group of Dr. David Mendoza-Cózatl (University of Missouri) for allowing us to use their Imaging PAM machine. Funding for this work provided by National Science Foundation, IOS-2414183 and IOS-2343815 (RM). CAS acknowledges financial support for this project from a University of Missouri College of Agriculture, Food and Natural Resources Joy of Discovery Grant Discovery.

## Notes

### Competing Interest Statement

The authors have declared no competing interest.

