## Supplementary Figures for "The proteotoxicity of azetidine 2-carboxylic acid is associated with reactive oxygen species accumulation"

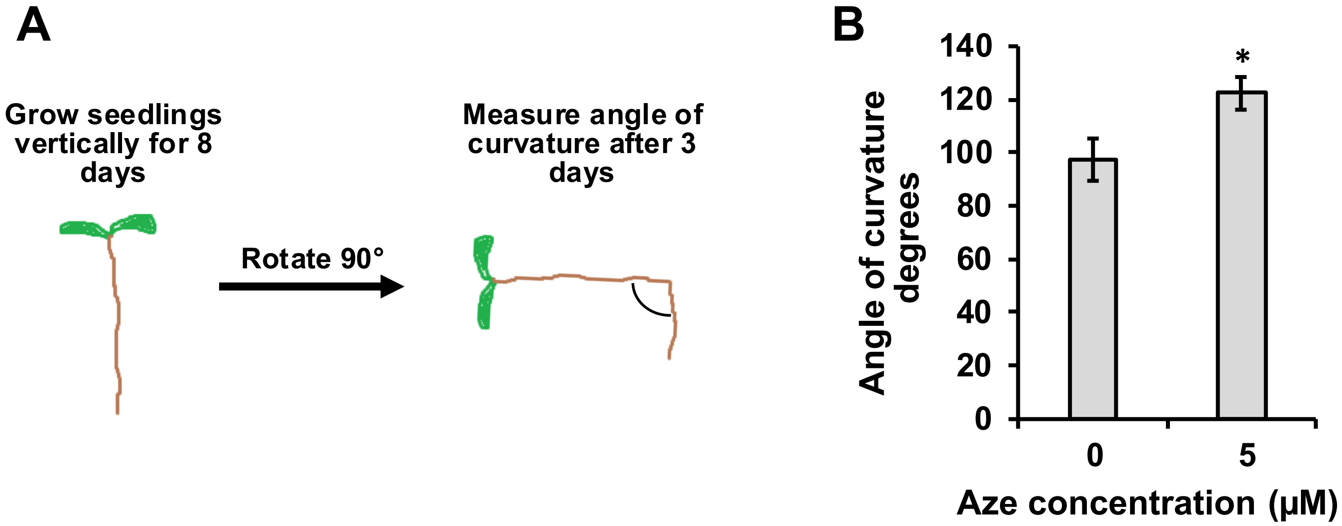


**Supplementary Fig. 1 | Aze alters root gravitropic response.** Arabidopsis was grown on 0.5x MS media for 8 days vertically supplemented with 0 µM or 5 µM Aze. Plates were rotated 90 degrees and after 3 days angle of root curvature was measured using imageJ. **(A)** Cartoon schematic of experimental setup, **(B)** average root curvature of the primary root following three days after rotation. Arabidopsis grown on 0 µM Aze showed a full gravitropic response of ~ 90°, whereas Arabidopsis grown on 5 µM had a reduced gravitropic response of ~ 120°. Bars indicate mean ± SEM of ≥ 10 biological replicates. Stars indicate significant differences from 0 µM Aze control in a one-way T-test of P < 0.05. Abbreviations: Aze: Azetidine 2-carboxylic acid.

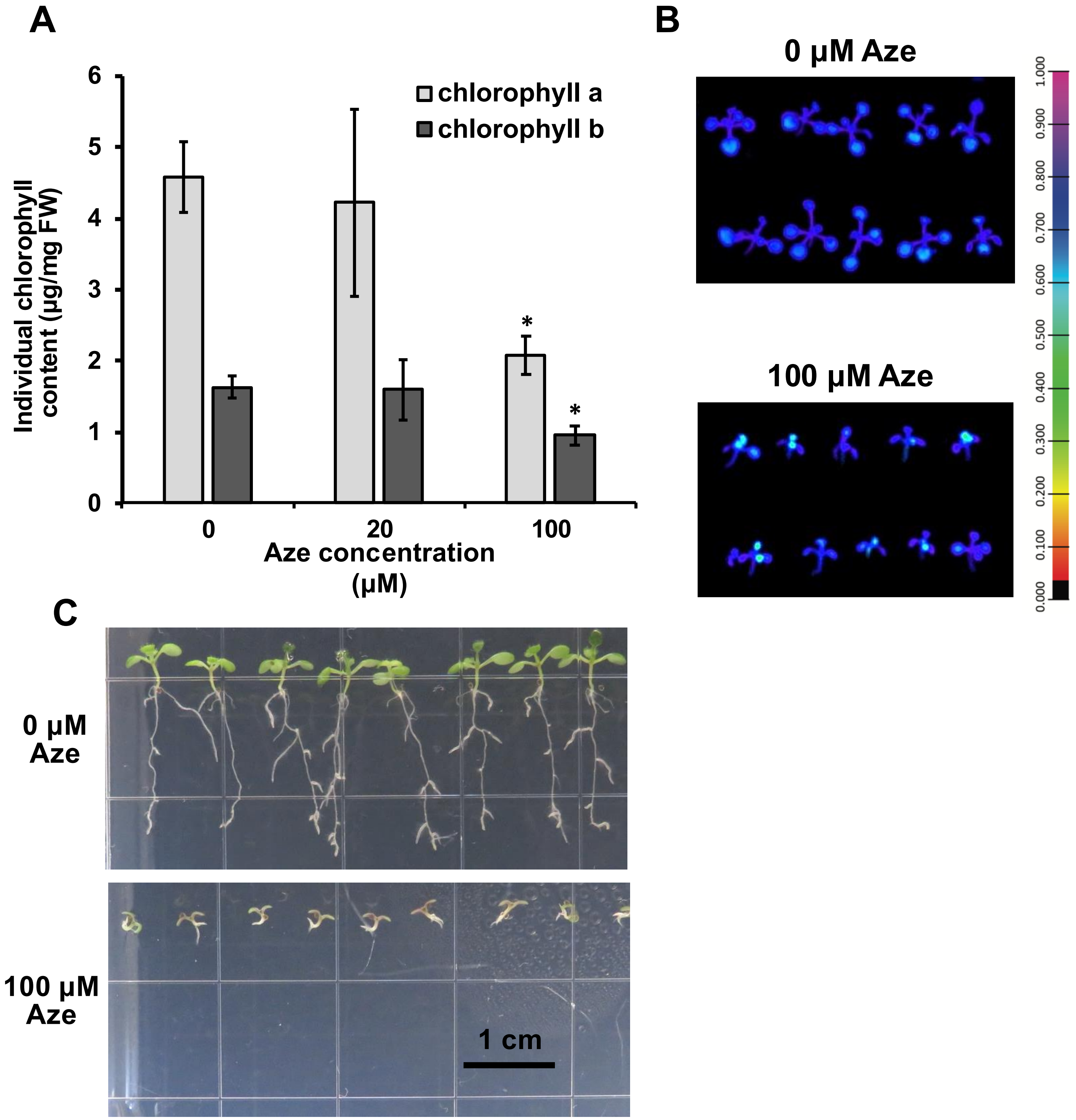


**Supplementary Fig. 2 | Aze alters photosynthetic capacity of Arabidopsis.** Arabidopsis was grown on 0.5x MS media and supplemented with varying amounts of Aze. Above ground stress phenotypes were measured: **(A)** chlorophyll content broken down into the individual chlorophyll compounds a and b. Stars indicate significant differences from 0 µM Aze control in a one-way T-test of $P < 0.05$. **(B)** A representative image of Fv/Fm determined following saturating light pulse used to quantify photosynthetic capacity. Old levels of Aze treated Arabidopsis have lower Fv/Fm. The Fv/Fm intensity scale is shown on the right. Quantification of images is shown in Fig. 2. **(C)** Arabidopsis were grown for 10 days on 0 or 100 µM Aze, showing visible purple pigment accumulation in plants grown on Aze. Abbreviations: Aze: Azetidine 2-carboxylic acid, FW: Fresh weight, Fv/Fm: maximum quantum efficiency of photosystem II.

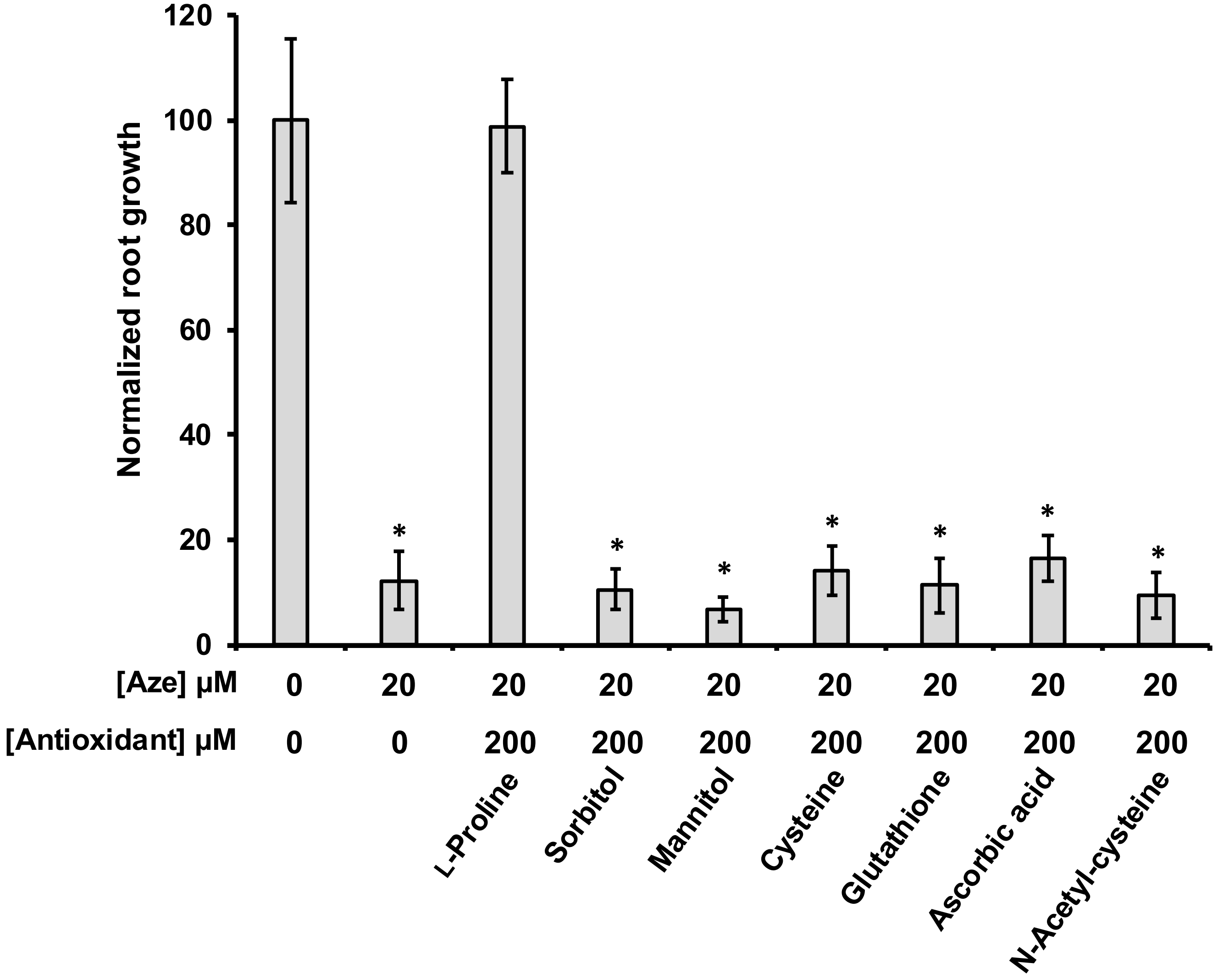


**Supplementary Fig. 3 | Aze induced root reduction is not restored by addition of antioxidants.** Arabidopsis was grown on 0.5x MS media for 8 days supplemented with 20 µM Aze and 200 µM of various antioxidants. Root growth was measured from images using imageJ. Root growth was normalized to 0 µM Aze. In this experiment Aze inhibited growth by around 85%, which was fully recovered by adding proline, however, none of the antioxidants tested were able to restore root growth. Bars indicate mean ± SEM of ≥ 10 biological replicates. Stars indicate significant differences from 0 µM Aze control in a one-way T-test of $P < 0.05$. Abbreviations: Aze: Azetidine 2-carboxylic acid.
